# Assay concordance sets exact ceilings on what one biological score can predict

**DOI:** 10.64898/2026.08.24.746774

**Authors:** Zongmin Liu

## Abstract

Computational models of biology are ranked by averaging one prediction against many experimental realizations of a phenotype that are treated as interchangeable. We show this imposes an exact, model-free ceiling fixed by how much those realizations agree with each other, and that the ceiling depends on the evaluation metric through a single support-function identity. Measuring assay concordance across four public registries—2,822 MaveDB score sets, 217 ProteinGym assays, two drug screens and 1,150 CRISPR cell lines—we find that two assays of one target agree at 0.56–0.68, and that 541 domains measured twice with different proteases fix assay reliability at 0.897, so 70–90% of every ceiling is irreducible biology rather than noise. Published predictors realize 63% of the achievable on the correlation benchmarks report and 18% on the top-1% selection their users perform. We provide the estimator, the ceilings, and the measurements the field has not made.

---

Progress in computational biology is largely read off scalar benchmark scores. ProteinGym ranks variant-effect predictors against 217 deep mutational scans ^1^, built on a decade of deep mutational scanning ^2–4^; single-cell foundation models are ranked on perturbation panels ^5^; drug-response models are scored across cell lines ^6,7^; gene-dependency models across CRISPR screens ^8,9^. In each case the reported quantity averages one prediction over experimental realizations of a phenotype, and progress is the change in that average.

The field cannot agree on what these averages mean. Deep perturbation models do not outperform linear baselines ^5^; protein language models stop improving with scale ^10^; single-cell pretraining corpora give inconsistent gains ^11,12^. These disagreements are usually attributed to benchmark construction or tuning—the same diagnosis reached repeatedly in machine learning ^13–16^—and the remedies proposed are better splits and better baselines.

We argue that a structural constraint has been overlooked. The experiments being averaged are not replicates of one latent quantity; they measure *different* quantities that share a name. A *β*-lactamase ampicillin-resistance scan and an amoxicillin-resistance scan are not two noisy views of “fitness”; an abundance assay and an enzymatic-activity assay of one protein measure different biology. Once that is taken seriously, a predictor emitting *one* score per item faces a hard geometric constraint—computable from assay data before any model is trained.

Previous benchmarking studies have documented imperfect inter-assay agreement and cautioned that experimental correlation itself constrains model–experiment agreement ^17–19^. A contemporaneous benchmark likewise estimates a ceiling from technical replicates and places frontier models at roughly 54% of it ^20^; target-dependent maxima for precision@*m* are also known in ranking ^21,22^. What has been missing is a general quantitative theory that turns assay concordance into the exact feasible set for arbitrary *k*, derives closed-form ceilings for the metrics a field actually reports or uses, separates measurement noise from biological discordance, predicts those ceilings out of sample, and identifies the training objective that reaches the relevant frontier. Those are the quantities we supply.

## The feasible set is an ellipsoid

Fix a target and *n* items assayed in *k* **contexts**—assays, readouts, cell backgrounds. Let *y*_*j*_ be the standardized measurement in context *j* and *R*_*jl*_ = ⟨*y*_*j*_, *y*_*l*_⟩*/n* the **concordance matrix**, a property of the experiments alone. A **context-agnostic** predictor emits one standardized score *s* used in every context, with performance profile *ρ*_*j*_ = corr(*s, y*_*j*_). This is the operative regime for zero-shot variant-effect predictors ^23–28^, for a perturbation embedding reused across cell contexts ^29^, and for any generalist model chosen off a leaderboard and applied downstream.

The Gram matrix of (*s, y*_1_, …, *y*_*k*_) must be positive semidefinite, and its generalized Schur complement ^30^ gives the exact feasible set (Theorem 1, Supplementary Note 1):

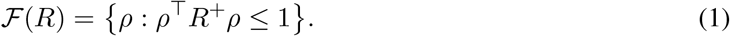

No assumption on model class, capacity, pretraining or compute enters. Scaling cannot leave the ellipsoid; only conditioning the score on the context can. Two closed forms follow. The best attainable mean is 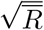 with 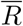 the mean of all *k*^2^ entries, attained by the context-averaged phenotype 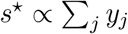 (Corollary 2); the best attainable *worst-context* value is (**1**^*⊤*^*R*^*−*1^**1**)^*−*1*/*2^ when *R*^*−*1^**1** ≥ 0 and a small convex program ^31^ otherwise (Corollary 3). For 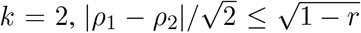, so a family of predictors with different inductive biases must spread along an axis whose length is set by assay discordance (Fig. 1a,b). The bound is a sharp partial-identification statement ^32^ rather than an approximation, and it binds: across all 3,417 model × assay-pair observations no published predictor lies outside the ellipse measured for its pair (Extended Data Fig. 1).

**Figure 1.**
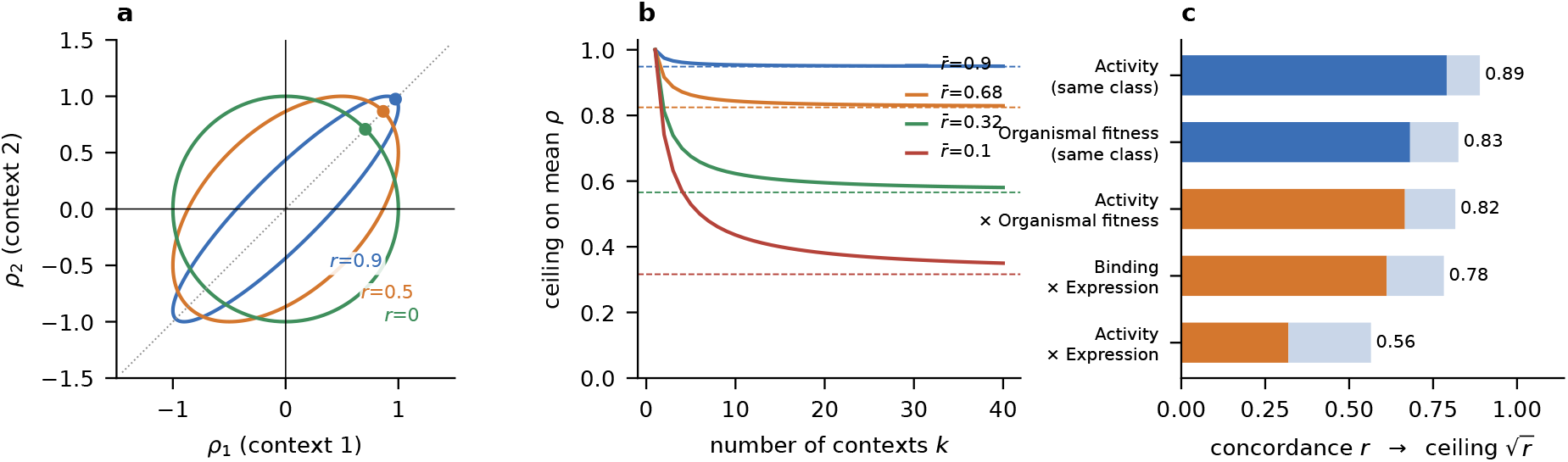
Assay concordance fixes an exact feasible set. **a**, The feasible set of performance profiles for a context-agnostic score over two contexts is an ellipse with semi-axes 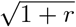 and 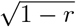; dots mark the optimum of the mean. **b**, Ceiling on the mean correlation against the number of contexts, for four values of mean concordance; dashed lines are the *k* → ∞ limits 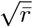, Concordances measured within ProteinGym targets and the ceilings they imply (light extension). Blue, two assays of one phenotype class; orange, two classes.

For *k* = 2 replicates Corollary 2 reduces to Spearman’s correction for attenuation ^33^, later extended to correlated errors ^34,35^ and to the multitrait–multimethod setting ^36^. What is new is the feasible *set* for arbitrary *k*, the closed forms over it, and the reading of the off-diagonal of *R* as biological discordance between non-exchangeable contexts rather than as unreliability—a limit that better instruments do not remove (Supplementary Note 2).

### One identity covers every metric

Correlation is what benchmarks report, not what anyone uses. A group choosing 96 variants to synthesize cares about the top of a ranking; a clinical classifier cares about AUROC, which is how functional evidence enters variant interpretation ^37–40^. Call a measure *representable* if *M* (*s, y*_*j*_) = ⟨*g*(*s*), *h*(*y*_*j*_)⟩, and let *G* be the achievable set of score representations. Then (Theorem 12)

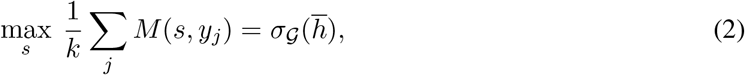

the support function of *G* at the context-averaged label 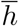. The content is in *G*: a **unit sphere** gives 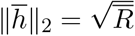, so Corollary 2 is the spherical case; the *m***-hot vectors** give the top-*m* ceiling (*km*)^*−*1^ Σ *c*_*i*_ over the *m* largest **context counts** *c*_*i*_ = #{*j* : *i* ∈ top-*m*(*y*_*j*_)}, attained by selecting the items appearing in the most contexts’ top lists; the **permutations** give the AUROC ceiling by the Mann–Whitney identity ^41^ and the rearrangement inequality ^42^. If the contexts’ top lists are disjoint the selection ceiling is 1*/k* while the correlation ceiling can stay high, so the two read different parts of the distribution and neither bounds the other. All are exact and computable in *O*(*n* log *n*).

### Two cohorts agree that concordance is far from one

MaveDB is the registry for multiplexed assays of variant effect ^43,44^, so two score sets on one gene are the comparison the theory needs. Of 557 published score sets with a mapped gene, 62 genes carry at least two; we retrieved 216. Score sets do not share a sign convention (104 of 295 raw pairs are negative), so we oriented them with two anchors that use no model output—nonsense variants must score below missense, and the score must correlate non-negatively with BLOSUM62—which agree in **148 of 148** score sets where both apply (Methods).

Two independent cohorts give overlapping ceilings, and the larger gives the lower value, so the ProteinGym figure is if anything optimistic (Fig. 2a, Supplementary Table 1). The 136 within-gene MaveDB pairs over 24 genes agree at 0.555, a ceiling of **0.745** [0.700, 0.791]; ProteinGym’s 18 same-class pairs over 7 proteins agree at 0.681, a ceiling of **0.825** [0.750, 0.867] (per gene in Extended Data Fig. 2 and Supplementary Table 2).

**Figure 2.**
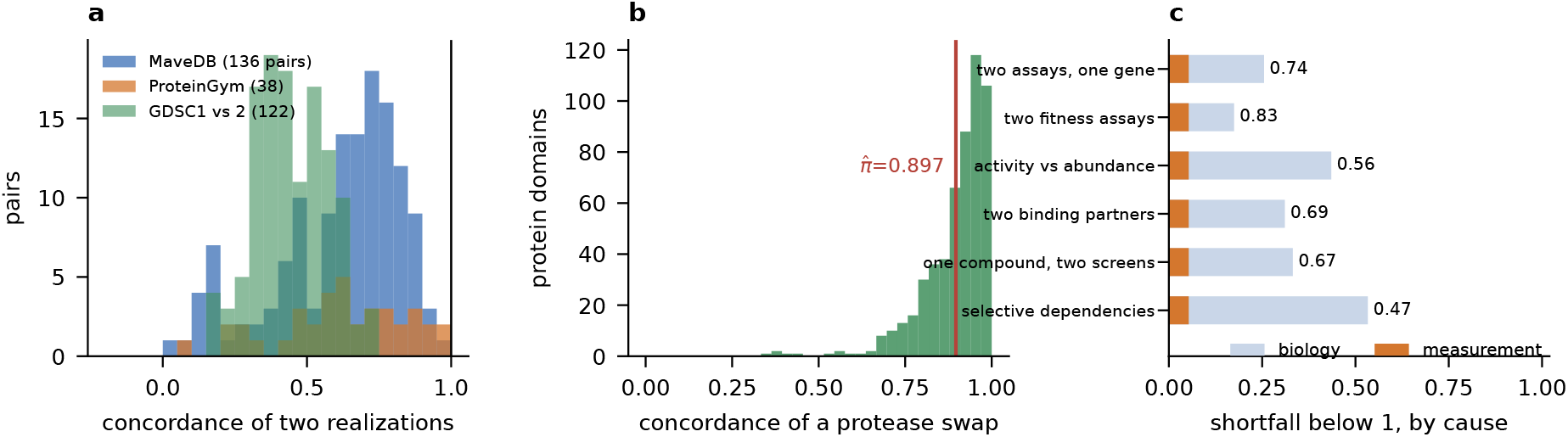
Two independent cohorts, and the split between noise and biology. **a**, Concordance between two realizations of a phenotype: within-gene pairs in MaveDB, within-protein pairs in ProteinGym, and the same compound in two drug screens. None approaches 1. **b**, Concordance of a protease swap—one domain, one library, one platform—across 541 domains, giving 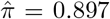. **c**, Each ceiling’s shortfall below 1, split into the part a protease swap accounts for (measurement) and the remainder (biology); labels give the ceiling.

Within a ProteinGym protein, assays of *different* phenotype classes agree much less: activity × organismal fitness 0.666, binding × expression 0.612, **activity** × **expression 0.319** [0.17, 0.51]. Activity and abundance assays of one protein are close to measuring different things. Intervals are cluster bootstraps ^45^ over targets widened 1.4× because simulation at these designs shows the raw interval covers only 84–87% (Supplementary Note 3). The best of 96 published predictors attains 0.491 against the fitness ceiling of 0.825 and 0.517 against the activity ceiling of 0.889—**59% and 58%**.

### Most of every ceiling is biology, not noise

A ceiling reflecting measurement error would be an engineering problem; one reflecting biological difference is not. Writing 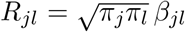 with *π* the reliability of an assay, *π* is unidentified without a replicate—and the public record contains one at scale. The mega-scale stability studies ^46^ deposit, for each of **541 protein domains**, two score sets from one library, platform and laboratory differing only in the protease reporting folding stability. These give 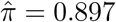 [0.888, 0.905] (median 986 shared variants per pair; 1% of domains below 0.5). Classifying every MaveDB pair by what actually differs between the two measurements yields the ordered reliability ladder required by the decomposition (Extended Data Fig. 3, Supplementary Note 4), and every ceiling then decomposes (Fig. 2b,c; Supplementary Table 3). Measurement accounts for 21% of the gap to 1 for two assays of one gene, 30% for two organismal-fitness assays, 12% for activity against abundance, 16% for one compound in two drug screens and 10% for selective gene dependencies.

#### Between 70% and 90% of every ceiling here is irreducible biological difference

Repeating the experiments more carefully would move the fitness ceiling from 0.825 to at most 0.872. A protease swap holds library and platform fixed but changes the reporter, so 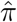 is a *lower* bound on reliability and the measurement shares *upper* bounds.

### The ceiling predicts out of sample

A bound estimated on the data it constrains is a description. Predicting each protein’s ceiling from the class concordance of *other* proteins only, the best model exceeded the prediction on **0 of 20** proteins and **0 of 1**,**900** model × protein observations, with mean signed error +0.009. The MaveDB ceiling of 0.745—a different registry, different genes, no predictor scores—is likewise exceeded by no model on any multi-assay protein. And the bound is informative rather than loose: across proteins it tracks attainment at *r* = +0.676 (permutation *P* = 0.0014, Extended Data Fig. 4).

### The field reports the metric on which it does best

The ceilings above are class-level; those below are per-target, averaged over the 19 proteins with at least 500 shared variants, which is why the correlation ceiling reads 0.866 rather than 0.825. Holding models, data and targets fixed and varying only the metric, the ceiling moves from 0.866 for the correlation the benchmarks report, through 0.950 for AUROC at 20% positives, to 0.626, 0.580 and **0.525** for selecting the top 10%, 5% and 1%; the best published model attains 0.550, 0.727, 0.248, 0.166 and 0.099 against them (Fig. 3a, Supplementary Table 4; every target in Extended Data Fig. 5).

**Figure 3.**
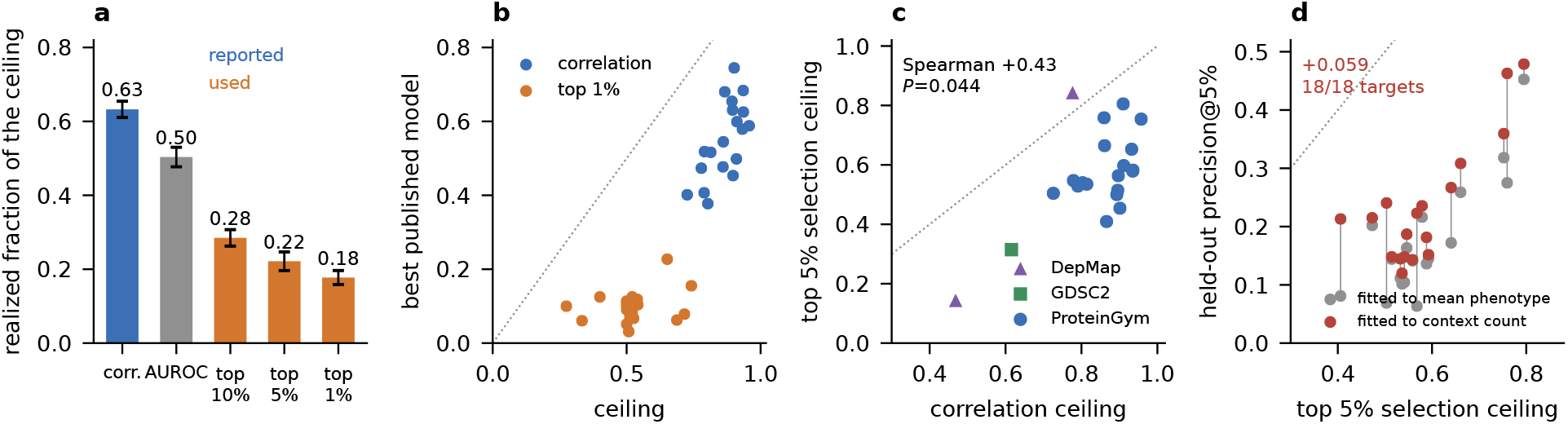
The ceiling depends on the metric, and the field reports the one it does best on. **a**, Realized fraction of the exact ceiling for the same 95 predictors on the same 19 proteins under five metrics; error bars are s.e. over proteins. **b**, Ceiling against best attained per protein, for correlation and top-1% selection. **c**, Correlation ceiling against top-5% selection ceiling across 22 targets in three domains. **d**, Held-out precision at the top 5% for the same feature bank fitted to the conventional mean-phenotype target and to the Corollary 13 context-count target, against the selection ceiling.

Realized fraction is (attained − chance)/(ceiling − chance): 0 at chance, 1 on the frontier, and comparable across metrics as raw scores are not. **The field has realized 63% of the achievable on the metric it reports and 18% on the selection task its users perform**—same models, assays and targets. The selection ceiling is itself lower (median 0.61 of the correlation ceiling at the top 1%) because contexts disagree most in the tail, but the attained value falls much further, so the shortfall is genuine (Fig. 3b). Across all 22 targets in three domains the two ceilings correlate at Spearman +0.43 (*P* = 0.044): related, not interchangeable (Fig. 3c). All confirmatory tests reported here are theory-specified and survive Holm correction ^47^ at family-wise *α* = 0.05.

The shortfall is partly a matter of optimising the wrong functional, and Corollary 13 says which one is right. Refitting the same 192-feature bank on the same folds to the *context count c*_*i*_ rather than to the mean phenotype raises held-out precision at the top 5% from 0.175 to **0.235**, improving **18 of 18 targets** (+0.059 ± 0.014, *t* = 4.11), and lifts the realized fraction from 0.22 to 0.33—a 53% relative gain that also beats the best published score (+0.062 ± 0.024, *t* = 2.54). At the top 1% the gain over the conventional target persists (+0.067 ± 0.022, *t* = 3.08) though it no longer separates from the best published score. Nothing changed but the training objective (Fig. 3d, Supplementary Table 5).

### Aggregate rankings are reproducible; per-context verdicts are not

Resampling which assay represents each multi-assay protein (1,000 draws) never changed the top-ranked model or the top-5 set; the induced s.d. of a model’s aggregate score is 0.0020, five times *smaller* than the item-level bootstrap error already reported. Restricted to one protein and one phenotype the picture inverts: the top-ranked model changes in **81%** of swaps between two assays of one protein that share a selection type, and pairwise reversal falls steeply with margin (46% at ≤ 0.005, 2% above 0.15; Supplementary Table 6), so a single-assay margin must exceed ≈ 0.15 to be reproducible while the published top ten spans 0.056 (Fig. 4a, Supplementary Note 5).

**Figure 4.**
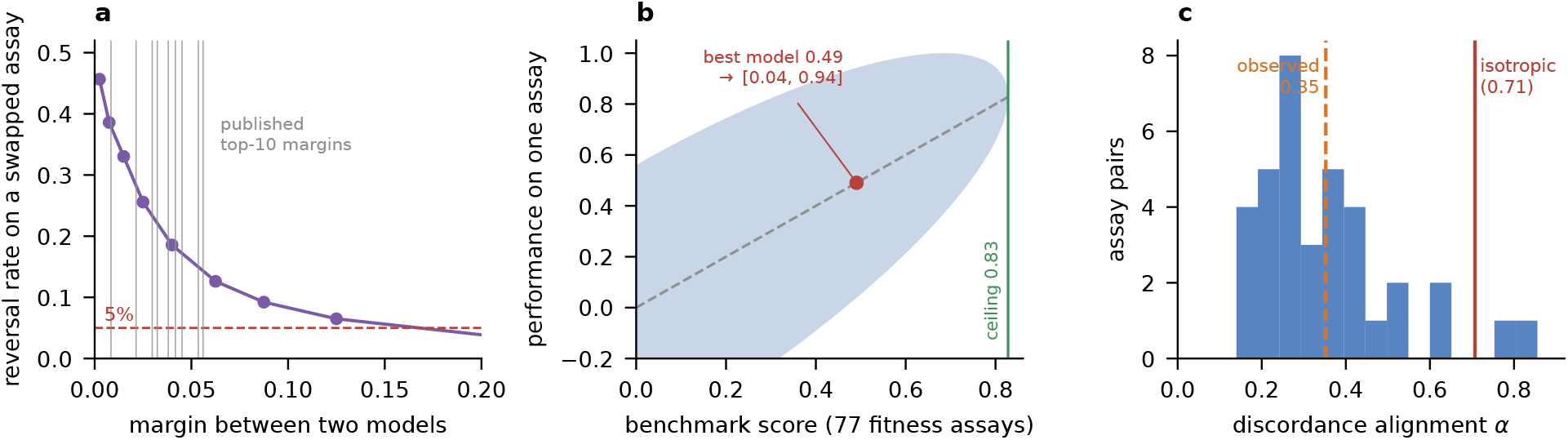
Aggregate reproducibility and per-context blindness have one cause. **a**, Probability that a pairwise model comparison reverses when the assay is swapped for another of the same protein and class, against the margin; grey lines are the nine published top-10 margins. **b**, Single-assay performances compatible with a given benchmark score over 77 organismal-fitness assays (Theorem 10); the best published model sits where the interval is [0.04, 0.94]. **c**, Discordance alignment 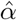 per assay pair: published models place 15% of their difference energy where assays disagree, against 0.707 for isotropic differences.

Theorem 10 reconciles these and converts a leaderboard number into a deployment interval. With *b* = *w*^*⊤*^*ρ* observed and *κ*_t_ = corr(*yt*, Σ_*j*_ *w*_*j*_ *y*_*j*_) the **benchmark–target concordance**, *ρ*_*t*_ is confined to a band of half-width 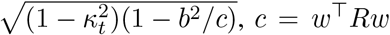. For an exchangeable class *κ* equals the ceiling itself, so a predictor scoring 0.49 across 77 fitness assays is pinned only to **[0.04, 0.94]** on any single one (Fig. 4b; by class in Extended Data Fig. 6). Both facts have one cause: decomposing model differences into the directions where assays agree and disagree, published predictors place only **15.2%** of their difference energy where assays disagree (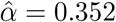 against 0.707 for isotropic differences). A benchmark stable *because every candidate shares a blind spot* has not shown the blind spot is small.

### The metadata suffices; the models do not use it

Selecting the best model per assay gains +0.073 Spearman, improving 165 of 165 proteins, so the interaction is real and large. Gating on recorded metadata gains +0.002 (n.s.), and a gradient-boosted gate on every field gains nothing. Theorem 8 shows this is a modelling limit, not missing information: a predictor told a descriptor is capped at 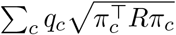, which for a hard partition is 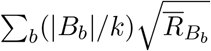, so a field’s worth is computable before a gate is trained and is monotone in Blackwell informativeness ^48^. ProteinGym’s selection type is worth +0.101 of a maximum +0.133 (MSA depth: +0.005), and the field realizes +0.002 of it (Fig. 5a, Extended Data Fig. 7). The closest prior benchmarking study handles multi-assay targets by keeping only the assay with the highest median correlation to predictors ^17,18^—the rule most likely to inflate apparent attainment.

Conditioning at the score level does recover part of it. Ridge recomposition of the 95 published scores on held-out variants gains +0.032 in 20/20 proteins, and—as the theory requires—the gain grows with discordance (*r* = +0.730, *P* = 2.6 × 10^*−*4^) and never exceeds its bound 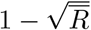. The reachable ceiling 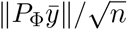 then separates a conditioning failure from an incomplete bank: adding a 44-feature read-out of ESM-2 650M ^49^ raises it on **20 of 20** proteins (+0.031, *t* = 11.8), and 53 hand-written biochemical features raise it *again* on top of that, also 20/20. A bank containing 16-billion-parameter retrieval-augmented models therefore fails to span a direction that a linear read-out of a mid-sized public model supplies. A gradient-boosted context-conditional head on all 192 features reaches 0.739 against the 0.866 ceiling—**85%**, from 64% for the best single published model (Fig. 5b, Supplementary Table 7).

**Figure 5.**
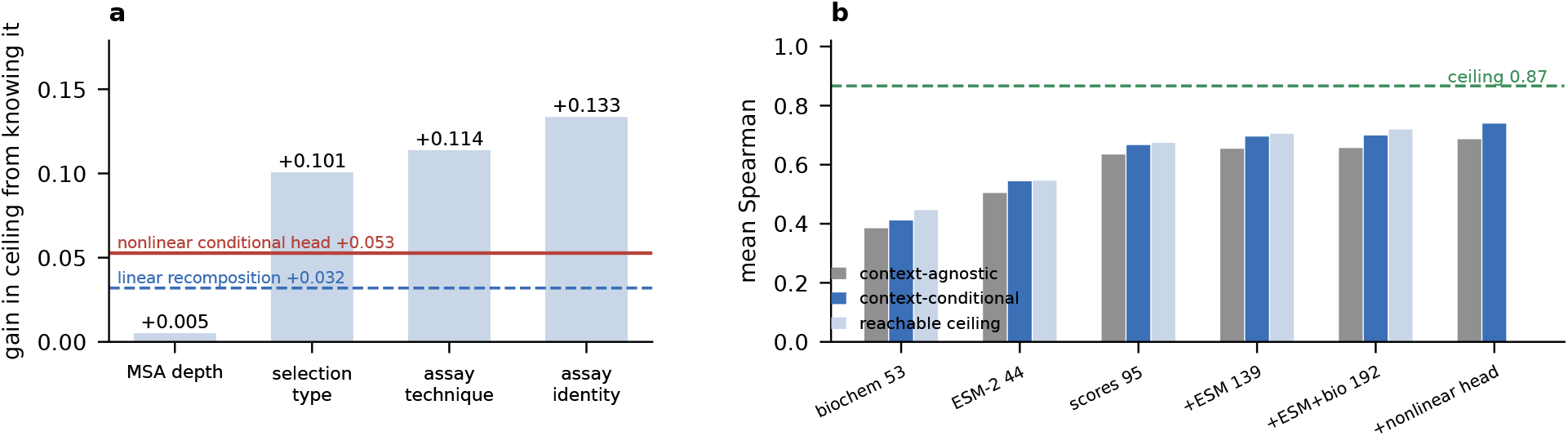
What a descriptor is worth, and what the predictor bank is missing. **a**, Exact gain in ceiling available from each recorded descriptor (Theorem 8), against what recomposition realizes. **b**, Context-agnostic fit, context-conditional fit and reachable ceiling for five feature banks; adding an ESM-2 read-out and, independently, hand-written biochemistry raises the reachable ceiling on 20 of 20 proteins.

### The limits transfer, and bind harder

In GDSC2^6,7^ (504 cell lines × 166 compounds) a compound-agnostic score is capped at 0.615 and at 0.400 on its worst compound, while a cell-line-agnostic score reaches 0.852 for free—which is why pooled (drug, cell line) evaluations report high correlations. GDSC1 reproduces the latter (0.861) and gives a *lower* compound-agnostic ceiling (0.483) because its panel spans 344 compounds rather than 166, exactly as Corollary 7 predicts. For the 122 compounds in both screens the same compound on the same cell lines agrees at *r* = 0.446 in `LN_IC50`, capping any predictor of “the” response at 0.668 (Fig. 6a). That is a ceiling on a readout, not on a phenotype: re-engineering the response metric raises cross-study agreement, as Theorem 12 predicts, which makes the drug domain the weakest of the three (Supplementary Table 3). The ceiling follows the compound mix, ranging by target pathway from 0.528 (apoptosis regulators) to 0.831 (EGFR signalling) (Extended Data Fig. 8, Supplementary Table 8).

**Figure 6.**
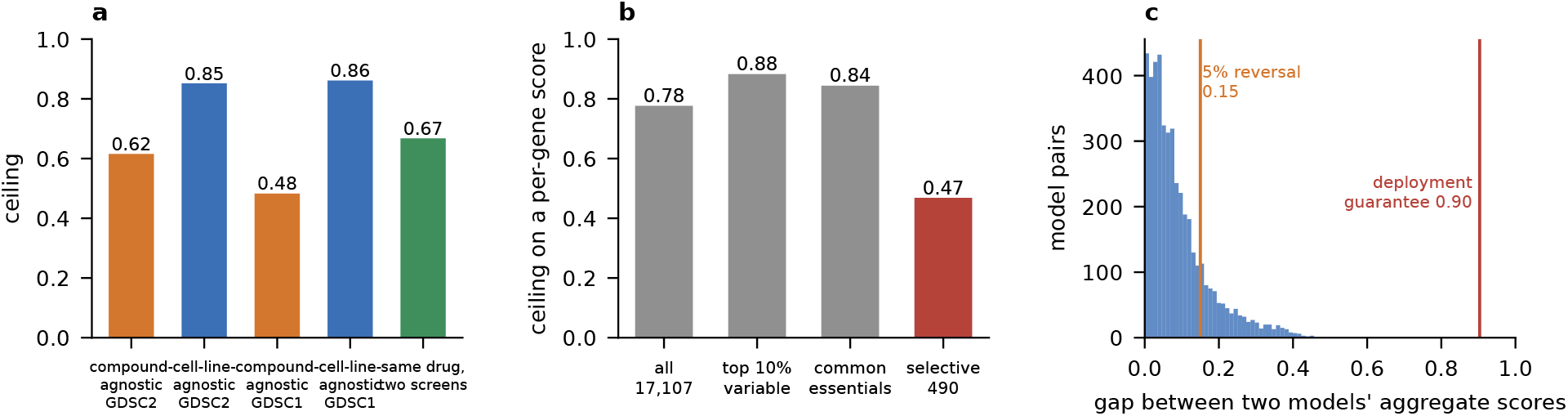
The limits transfer to two further domains and change what the numbers mean. **a**, Compound-agnostic and cell-line-agnostic ceilings in GDSC2, with the GDSC1 replication and the cross-study replicate anchor. **b**, Ceiling on a per-gene essentiality score in DepMap over all 17,107 genes and over subsets; it collapses to 0.47 on the 490 selectively essential genes. **c**, Gaps between the aggregate scores of 4,560 model pairs, with the empirical 5%-reversal threshold and the deployment guarantee marked.

In DepMap ^8,9^ (1,150 cell lines × 17,107 genes) a per-gene essentiality score reaches 0.777 over all genes—almost all of it free, since most genes are non-essential everywhere. Filtering to the most variable genes *raises* the ceiling to 0.88, because variance is dominated by the consistent essential/non-essential split. Only excluding common essentials, leaving the **490 selectively essential genes** that are candidate targets, does concordance collapse to 0.218 and the ceiling to **0.468** (Fig. 6b). The standard variance filter selects the easy cases.

### What changes downstream

Against the aggregate bootstrap, 3,879 of 4,560 model pairs (85%) are separable; against the Theorem 10 deployment guarantee, none is—an 18-fold mismatch (Fig. 6c). Across the 26 predictors stating a parameter count, size does not predict aggregate performance (*r* = +0.19, *P* = 0.36) but predicts *less* behavioural diversity (*r* = −0.568, *P* = 0.002): scaling has been buying similarity. The axis they spread along is interpretable: structure-based inverse-folding models at one end, autoregressive sequence models at the other (Supplementary Table 9). Observational on 26 models, but a mechanistic account of 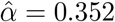 (Extended Data Fig. 9).

## Discussion

Six numbers become reportable, each computable from assay data before a model exists: the ceiling 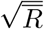 the profile rather than the scalar; the missing concordances (no protein has been assayed for both in-vitro stability and organismal fitness at variant scale, Supplementary Table 10); the discordance alignment 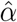, which says whether a leaderboard’s stability is earned; the deployment interval *κ* and *H*; and the reachable ceiling, which says whether a field’s collective output spans the phenotype it predicts. Ours does not.

Theorem 12 makes several projects well posed, one of which we settled here: because the optimal context-agnostic selection is the set with the largest context counts, retargeting an existing feature bank at that objective improves held-out selection on 18 of 18 proteins. Two others remain open. The distribution of context counts is a new statistic of an assay panel, with its own estimator and biological interpretation. And registries depositing per-replicate scores would make the noise/biology split identifiable for every assay: ours was recoverable only through the accident of the protease pairs, and reporting standards for these assays already exist ^50^ to extend.

The framework has three defined scopes of interpretation. Theorem 1 bounds one score serving several contexts on the *same items*, not a benchmark average over targets that each carry a single context. The ellipsoid audit tests the context-agnostic regime; exact score agreement across assay files establishes that regime directly for the published predictors analysed here. Class-level ceilings treat two assays of one class on one target as exchangeable realizations of that phenotype; a within-protocol replicate series would permit this assumption to be tested directly. Additional scope conditions and sensitivity analyses are detailed in Supplementary Note 6.

## Data availability

All data are public. ProteinGym v1.3 (MIT): https://marks.hms.harvard.edu/proteingym/, Zenodo 10.5281/zenodo.15293562. MaveDB: 2,822 published score sets via https://api.mavedb.org/api/v1/. GDSC1 and GDSC2 release 8.4: https://ftp.sanger.ac.uk/pub/project/ cancerrxgene/releases/. DepMap 24Q2 Public `CRISPRGeneEffect.csv`: figshare article 25880521. ESM-2 650M weights: https://dl.fbaipublicfiles.com/fair-esm/. Source data are provided for every main and Extended Data figure.

## Code availability

`ceiling.py` is a standalone numpy-only implementation of every estimator reported here. All analyses, five numerical verification suites for the fourteen theoretical results, and an audit script that re-derives every number quoted here from the saved data files are included. To be deposited at Zenodo with a persistent DOI on acceptance.

## Author contributions

Z.L. conceived the study, developed the theory, performed all analyses and wrote the manuscript.

## Acknowledgements

We thank the ProteinGym, MaveDB, GDSC and DepMap consortia for making their measurements and baseline predictions openly available; this work exists only because they did.

## Methods

### Theory and its numerical verification

All fourteen theoretical results are stated and proved in Supplementary Note 1. Each is additionally checked against brute-force optimization, which is the practice we would want applied to any bound of this kind.

Theorem 1 (feasible set) is verified by explicitly constructing, for randomly drawn target profiles inside the ellipsoid, a score vector realizing them: 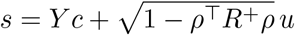 with *c* = *R*^+^*ρ* and *u* orthogonal to the column space of *Y* ; maximum error over 300 random designs 7 × 10^*−*16^. Corollary 2 is checked against numerical maximization of the mean over unit-norm score vectors (maximum error 9 × 10^*−*7^). Corollary 3 is non-smooth in its raw form, so it is checked by solving the smooth epigraph program over the ellipsoid with SLSQP from 12 restarts (maximum error 3 × 10^*−*12^); the hypothesis *R*^*−*1^**1** ≥ 0 is separately shown to be necessary by counterexample search over random correlation matrices. Corollary 4 is checked by violation search over 2 × 10^4^ random score vectors; note that the bound requires 1 + |*r*|, since for *r <* 0 the feasible point 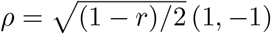 violates the familiar 1 + *r* form. Every assay pair measured in this work has *r >* 0 (minimum 0.054), so the two forms coincide on our data. Corollary 7 is checked by monotonicity in *k* and *m* and against its *k* → ∞ limit.

Theorem 8 (partial information) is checked against direct maximization over one score vector per signal value, using random partitions of the context set: maximum excess of brute force over the closed form 0.0 across 60 designs. Its three specializations are checked against their closed forms (maximum error 4 × 10^*−*16^), and Corollary 9 (Blackwell monotonicity) by 400 refine/coarsen pairs with zero violations. Theorem 10 (resolution) is checked against constrained optimization on the ellipsoid slice over 300 random designs (maximum excess 8 × 10^*−*13^), including the degenerate *κ* = 1 case and the vanishing of the interval as 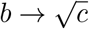. Theorem 12 and its corollaries are checked **exhaustively**: Corollary 13 against every *m*-subset of *n* items over 400 designs (maximum excess 1 × 10^*−*16^) and Corollary 14 against all *n*! rankings over 150 designs (maximum excess 6 × 10^*−*16^). We also construct a case where the correlation ceiling is 0.334 while the top-*m* ceiling collapses to 1*/k*, confirming that neither bounds the other.

### Simulation study of the estimator

Because the theory is exact, what can fail is estimation. On data generated from known ground truth (`code/simulation.py`) we established the following.

Results are in Supplementary Note 3 and Extended Data Fig. 10.

#### Bias and variance

The plug-in ceiling 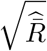 is essentially unbiased for the *Spearman* ceiling at every design tested (*n* ∈ {200, 10^3^, 5 × 10^3^}, *k* ∈ {2, 5, 20}, *r* ∈ {0.30, 0.68}; worst |bias| 0.0025, mean 0.0007), and its standard deviation falls as *n*^*−*1*/*2^ (0.0172 at *n* = 200 to 0.0034 at *n* = 5000, ratio 5.1 against 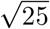). A Gaussian Pearson concordance of 0.68 corresponds to a Spearman concordance of 0.663 via 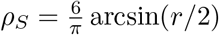, so mixing the two would understate the ceiling by up to 0.011. We use Spearman throughout, matching what the benchmarks report, and the ceiling we quote is therefore a Spearman ceiling.

#### Interval coverage and calibration

With protein-level heterogeneity in the true concordance (s.d. 0.12), the cluster bootstrap undercovers at small numbers of targets: 80.5% at 5 targets, 87.3% at 7 (the ProteinGym organismal-fitness design), 89.3% at 15, 95.0% at 40. A search over symmetric inflation factors applied to the bootstrap half-widths gives 1.4 for nominal 95% coverage at 7 targets (83.6% → 92.6% → 95.0% at factors 1.0, 1.2, 1.4). All intervals in the paper are widened by this factor and the raw intervals are also reported.

#### Power of the ellipsoid audit

For a partially context-aware predictor *s*_*j*_ = (1 − *ω*)*s*_0_ + *ωy*_*j*_, the audit has a false-positive rate of 0.000 at *ω* = 0, power 0.000 for *ω* ≤ 0.2, and power 0.80–1.00 at *ω* = 0.4 across *n* ∈ {300, 10^3^, 4 × 10^3^} and *r* ∈ {0.35, 0.70}. Zero violations therefore licenses “no published predictor is strongly context-aware”, not “every predictor is exactly context-agnostic”; the exact agreement of each model’s scores across assay files (median Pearson 1.0000) establishes the stronger statement directly.

#### Robustness

With reliabilities drawn with s.d. up to 0.35 the plug-in estimator continues to report the ceiling implied by the realized *R*; what changes is that the true ceiling itself falls below the equal-reliability value (by 0.036 at the largest spread). Heterogeneity costs interpretability, not validity, which is why the worst-context ceiling is also reported.

### Data

#### ProteinGym

Version 1.3 (MIT licence; Zenodo 10.5281/zenodo.15293562). We used the 217 substitution assays, the per-assay Spearman matrix for all released baselines, the reference file for UniProt identifier, taxon, sequence length, MSA length and selection type, and the per-variant scores of all released predictors. Of 103 score columns, 96 are complete across all 217 assays and 95 across all multi-assay proteins; only complete columns were used, so no comparison relies on imputation. Per-variant score files were retrieved member-by-member from the released archive by HTTP range request against the zip central directory.

#### MaveDB

Two cohorts. For the primary within-gene cohort, score sets were enumerated from `GET /api/v1/score-sets/mapped-genes` (557 sets with an HGNC mapping) and grouped by symbol; the 62 genes with ≥ 2 sets yielded 225 sets, of which 216 downloaded and parsed. For the reliability rung, that endpoint is insufficient because it excludes the designed and bacterial domains of the mega-scale stability studies, so we walked the experiment-set URN space (1–1400) via `GET /api/v1/experiment-sets/{urn}`, discovering 2,822 published score sets across 2,042 experiments and 1,273 experiment sets. Scores come from `GET /api/v1/score-sets/{urn}/scores`; variants are matched on protein-level HGVS where available (214 of 216 sets in the primary cohort), otherwise nucleotide-level, and pairs mixing notations are dropped. Duplicate variant rows are averaged.

#### GDSC

GDSC1 and GDSC2 fitted dose–response, release 8.4 (24 July 2022), `LN_IC50`. Complete blocks are formed by restricting to rows and columns with *>* 80% coverage and then dropping any remaining incomplete row or column, giving 504 × 166 (GDSC2) and 132 × 344 (GDSC1). Cross-study concordance uses compounds present in both screens with ≥ 100 shared cell lines, matched on compound and cell-line name, with replicate measurements averaged.

#### DepMap

DepMap 24Q2 Public `CRISPRGeneEffect.csv` (figshare article 25880521). Gene symbols are taken from column headers before the parenthesised Entrez identifier; the complete block retains genes with *>* 98% coverage and then drops any remaining incomplete row or column, giving 1,150 cell lines × 17,107 genes. Concordance matrices over cell lines use a random subsample of 400 cell lines for tractability, which affects sampling error and not the estimand. Gene subsets are defined by the across-line standard deviation and mean of the gene-effect score, with “common essential” taken as mean effect *<* −0.5 following DepMap’s convention.

### Concordance estimation

For each pair of assays on the same target we restricted to single substitutions present in both, averaged duplicate variant entries, and computed Spearman correlation on the shared set (median 2,654 shared variants in ProteinGym, 952 in the MaveDB primary cohort, 986 in the reliability rung). Pairs with fewer than 100 shared variants were dropped; the falsification test additionally required 100 and the recomposition experiments 300. Class-level concordance is the mean over pairs of the given class combination. Uncertainty is a nonparametric cluster bootstrap resampling *targets*, because assays on one target are not independent, with 4,000–8,000 replicates and the calibration factor above.

#### Orientation

MaveDB score sets do not share a sign convention: some report a functional score, others a depletion or damage score, and 104 of 295 raw pairs in the primary cohort are consequently negative. Sign is metadata rather than biology, so we place every score set on a common scale using two anchors that use no model output. The *nonsense anchor* requires that nonsense variants score below missense ones, and applies to the 148 sets carrying ≥ 5 nonsense and ≥ 20 missense variants. The *BLOSUM anchor* requires non-negative correlation with the BLOSUM62 substitution score across missense variants, and applies to 179 sets. Where both apply they agree in 148 of 148 cases. The primary analysis uses only pairs in which both sets carry the nonsense anchor, so no circularity is possible; three sensitivity analyses (BLOSUM orientation, |*r*| over all pairs, unorientable sets left unflipped) are reported in Supplementary Table 5. Targets are weighted equally in pooled estimates.

#### The reliability rung

Of the 2,042 discovered experiments, 541 contain exactly two score sets whose titles differ only in protease for the same domain; all 1,082 downloaded and parsed. 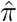 is the unweighted mean over domains with a percentile bootstrap over domains. MaveDB experiment membership alone does not define technical replication: score sets within one experiment can correspond to different proteases or to a protein measured with different binding partners. We therefore define the reliability rung only from the 541 matched domain pairs whose titles differ solely by protease, holding library, platform and laboratory fixed. Supplementary Note 4 reports the full measurement ladder.

### The ellipsoid audit

For each assay pair and each predictor we took the predictor’s score vector on the shared variants from one assay file, rank-normalized it once, and computed its Spearman correlation with each of the two measurement vectors, yielding (*ρ*_1_, *ρ*_2_) from a *single* score vector as Corollary 4 requires. Agreement between the two files’ copies of a predictor’s scores was verified (median Pearson 1.0000). We then evaluated the slack 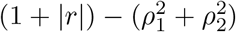 and the ellipsoid radius 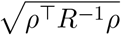 over 3,417 model × pair observations on 36 pairs and 21 proteins.

### Out-of-sample validation

For each multi-assay protein, the class-level concordance was re-estimated using only assay pairs from *other* proteins, with a global-mean fallback for empty cells, and the resulting predicted ceiling compared with what the 95 predictors attain. The cross-cohort test uses the MaveDB ceiling, estimated from a different registry with no predictor scores involved, against ProteinGym models. Informativeness is assessed by the correlation between a target’s measured ceiling and the best model’s performance on it, with a permutation null over 20,000 draws.

### Metric-general ceilings

Correlation ceilings use rank-standardized vectors throughout. Top-*m* ceilings apply Corollary 13 with *m* = ⌈*fn*⌉ for *f* ∈ {0.01, 0.05, 0.10} and a floor of 10 items, taking the top of each context by measured value; attainment is the mean precision@*m* of the model’s own top-*m* set against each context’s. AUROC ceilings binarise each context at its top 20% as the positive class, matching the sign convention of published variant-effect scores, and apply the Corollary 14 rearrangement bound; attained AUROC is computed in both orientations and the larger taken, so that a sign convention cannot make a competent predictor appear sub-chance. The realized fraction is (attained − chance)*/*(ceiling − chance) with chance = 0 for correlation, *m/n* for top-*m* and 0.5 for AUROC. Only proteins with ≥ 500 shared variants enter, giving 19 of the 20.

### Gating, recomposition and feature banks

All gating rules are fitted out of fold. Stratified gates select, for each held-out assay, the model with the highest mean out-of-fold score within the stratum defined by the listed metadata fields, falling back to the global winner when a stratum holds fewer than five out-of-fold assays; folds are leave-one-protein-out over 186 proteins. The learned gate is a histogram gradient-boosting regressor on (selection type, MSA-depth category, taxon, sequence length, MSA length, number of mutants, model identity) predicting per-assay Spearman, with 10-fold cross-validation grouped by protein. Inference is a paired comparison with the protein as the unit of independence.

For recomposition, scores and measurements were rank-normalized on the shared set of each protein with ≥ 2 assays and ≥ 300 shared variants. Variants were split at random into halves (20 repetitions), ridge coefficients estimated on the training half with the penalty chosen on a nested split from *λ* ∈ [10^*−*2^, 10^3.5^], and both fits scored on the held-out half where the ceiling is recomputed. The context-agnostic fit regresses the mean standardized measurement; the context-conditional fit regresses each context separately. The reachable ceiling is 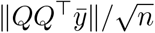 with *Q* an orthonormal basis of the held-out score matrix.

The biochemistry bank is 53 per-variant features: wild-type and mutant residue one-hots, BLOSUM62 score, signed and absolute changes in Kyte–Doolittle hydropathy, residue volume, formal charge and Pace– Scholtz helix propensity, relative sequence position and its distance to the nearer terminus, and indicators for mutant proline, wild-type glycine, charged mutant and buried-type wild type. The representation bank is 44 features: within each protein, the 24 leading principal components of the ESM-2 650M (`esm2_t33_650M_UR50D`) layer-33 residue representation of the wild-type sequence at the mutated position, plus the mutant-residue one-hot. Sequences longer than 1,022 residues were embedded in non-overlapping windows. No fine-tuning, supervision or MSA was used; embeddings were computed once on one GPU in under two minutes for all 23 targets. The nonlinear head is histogram gradient boosting (max depth 4, 200 iterations, learning rate 0.06, L2 = 1.0, early stopping on a 15% validation split), six random splits per protein, with identical splits, bank and evaluation as the ridge protocol.

### Selection-optimal training

Optimising ranking accuracy at the top of a list is a studied problem in its own right^49^; what Corollary 13 adds is the identification of the correct *target* under context heterogeneity. To test whether that objective helps, the same 192-feature bank was refitted on the same random half-splits (10 repetitions, targets with ≥ 800 shared variants, 18 of 20) to three different targets: the mean standardized phenotype, which is the conventional objective; the training-half context count *c*_*i*_, which Corollary 13 identifies as optimal for selection; and each context’s own top-*m* indicator separately, with the resulting scores summed. The ridge penalty is chosen on a nested split by held-out precision@*m* rather than by correlation, so each method is tuned for the criterion it is evaluated on. Evaluation is held-out precision@*m* against each context’s own top-*m* set with the Corollary 13 ceiling recomputed on the held-out half, and *m* is recomputed within each split so that the fraction is preserved.

### Descriptor value, resolution and consequences

Partition ceilings use Theorem 8 with posteriors uniform on each block of the partition induced by the metadata field, evaluated on the measured per-protein *R*. Theorem 10 intervals use uniform benchmark weights; at the class level *R* is the equicorrelated matrix implied by the measured within-class concordance and the class’s assay count, and *b* is the best published model’s class mean. Within proteins, *R* is the measured concordance matrix on shared variants.

The aggregate bootstrap resamples proteins (2,000 replicates) and a model pair is called separable when the paired difference excludes zero at 95%. Deployment separability requires the Theorem 10 intervals of the two models to be disjoint, i.e. a gap exceeding 2*H* evaluated at the organismal-fitness class. Parameter counts are parsed from model names by the pattern `<number><M|B>`; 26 of 96 names carry one. The general-skill and specialization coordinates are the leading right singular vectors of the class × model matrix before and after removing its rank-one component, with the general axis oriented so that larger means better. The full-mix deployment ceiling maximizes and minimizes *w*^*⊤*^*Mw* over the eight unmeasured entries of the class concordance matrix subject to entries in [−1, 1] and *λ*_min_(*M*) ≥ 0, by SLSQP from 30 random starts.

### Multiplicity and pre-specification

The theory fixes which comparisons matter before any data are seen, and we report every one we ran. The confirmatory tests are: no observation outside the ellipsoid; the context-conditional gain is positive; that gain grows with discordance; the gain never exceeds 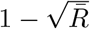; no model exceeds an out-of-sample predicted ceiling; and the ceiling correlates with attainment. Six directional tests, all specified by the theory rather than chosen after inspection; a Holm correction at family-wise *α* = 0.05 leaves every one significant, the largest *P* among them being 0.0014. Descriptive quantities — concordances, ceilings, variance decomposition, alignment, descriptor values, bank comparisons — are estimates with intervals and carry no *P* values. Analyses we regard as exploratory are labelled as such: the activity-class ceiling (3 pairs), the class-level completion bounds, the per-pathway GDSC ceilings, and the size-versus-diversity relation.

### Reproducibility

`code/audit.py` re-derives all 294 numbers quoted in this manuscript and its supplementary information from the saved data files and fails if any disagrees. `code/test_ceiling.py` independently reproduces every per-target ceiling, descriptor value, top-*m* ceiling and AUROC ceiling through the public API of `ceiling.py`, giving zero mismatches over 19–20 targets. Five numerical verification suites cover the fourteen theoretical results. Supplementary Note 7 records the independent audit checks applied to theorem statements, pair accounting, variance decomposition, interval calibration and AUROC orientation.

## References

[1] Notin, P. et al. ProteinGym: large-scale benchmarks for protein fitness prediction and design. Adv. Neural Inf. Process. Syst. 36 (2023).

[2] Fowler, D. M. & Fields, S. Deep mutational scanning: a new style of protein science. Nat. Methods 11, 801–807 (2014).

[3] Rubin, A. F. et al. A statistical framework for analyzing deep mutational scanning data. Genome Biol. 18, 150 (2017).

[4] Çubuk, H., Jin, X., Phipson, B., Marsh, J. A. & Rubin, A. F. Variant scoring tools for deep mutational scanning. Mol. Syst. Biol. 21, 1293–1305 (2025).

[5] Ahlmann-Eltze, C., Huber, W. & Anders, S. Deep-learning-based gene perturbation effect prediction does not yet outperform simple linear baselines. Nat. Methods 22, 1657–1661 (2025).

[6] Iorio, F. et al. A landscape of pharmacogenomic interactions in cancer. Cell 166, 740–754 (2016).

[7] Yang, W. et al. Genomics of Drug Sensitivity in Cancer (GDSC). Nucleic Acids Res. 41, D955–D961 (2013).

[8] Tsherniak, A. et al. Defining a cancer dependency map. Cell 170, 564–576 (2017).

[9] Dempster, J. M. et al. Chronos: a cell population dynamics model of CRISPR experiments. Genome Biol. 22, 343 (2021).

[10] Hou, C., Liu, D., Zafar, A. et al. Understanding language model scaling for protein fitness prediction. Nat. Comput. Sci. 6, 778–788 (2026).

[11] DenAdel, A. et al. Evaluating the role of pretraining dataset size and diversity on single-cell foundation model performance. Nat. Methods 23, 1447–1457 (2026).

[12] Kędzierska, K. Z., Crawford, L., Amini, A. P. & Lu, A. X. Assessing the limits of zero-shot foundation models in single-cell biology. bioRxiv (2023).

[13] Recht, B., Roelofs, R., Schmidt, L. & Shankar, V. Do ImageNet classifiers generalize to ImageNet? Proc. ICML (2019).

[14] Bowman, S. R. & Dahl, G. E. What will it take to fix benchmarking in natural language understanding? Proc. NAACL (2021).

[15] Liao, T., Taori, R., Raji, D. & Schmidt, L. Are we learning yet? A meta review of evaluation failures across machine learning. Adv. Neural Inf. Process. Syst., Datasets and Benchmarks Track (2021).

[16] Varoquaux, G. & Colliot, O. Evaluating machine learning models and their diagnostic value. Neuromethods 197, 601–630 (2023).

[17] Livesey, B. J. & Marsh, J. A. Variant effect predictor correlation with functional assays is reflective of clinical classification performance. Genome Biol. 26, 98 (2025).

[18] Livesey, B. J. & Marsh, J. A. Using deep mutational scanning to benchmark variant effect predictors and identify disease mutations. Mol. Syst. Biol. 16, e9380 (2020).

[19] Livesey, B. J. & Marsh, J. A. Updated benchmarking of variant effect predictors using deep mutational scanning. Mol. Syst. Biol. 19, e11474 (2023).

[20] De Brouwer, E. et al. AssayBench: an assay-level virtual cell benchmark for LLMs and agents. arXiv 2605.10876 (2026).

[21] Nemhauser, G. L., Wolsey, L. A. & Fisher, M. L. An analysis of approximations for maximizing submodular set functions. Math. Program. 14, 265–294 (1978).

[22] Fagin, R., Kumar, R. & Sivakumar, D. Comparing top k lists. SIAM J. Discrete Math. 17, 134–160 (2003).

[23] Frazer, J. et al. Disease variant prediction with deep generative models of evolutionary data. Nature 599, 91–95 (2021).

[24] Notin, P. et al. Tranception: protein fitness prediction with autoregressive transformers and inference-time retrieval. Proc. ICML (2022).

[25] Cheng, J. et al. Accurate proteome-wide missense variant effect prediction with AlphaMissense. Science 381, eadg7492 (2023).

[26] Brandes, N. et al. Genome-wide prediction of disease variant effects with a deep protein language model. Nat. Genet. 55, 1512–1522 (2023).

[27] Dauparas, J. et al. Robust deep learning-based protein sequence design using ProteinMPNN. Science 378, 49–56 (2022).

[28] Hsu, C. et al. Learning inverse folding from millions of predicted structures. Proc. ICML (2022).

[29] Replogle, J. M. et al. Mapping information-rich genotype–phenotype landscapes with genome-scale Perturb-seq. Cell 185, 2559–2575 (2022).

[30] Horn, R. A. & Johnson, C. R. Matrix Analysis 2nd edn (Cambridge Univ. Press, 2012).

[31] Boyd, S., Cortes, C., Mohri, M. & Radovanovic, A. Accuracy at the top. Adv. Neural Inf. Process. Syst. 25, 953–961 (2012).

[32] Manski, C. F. Partial Identification of Probability Distributions (Springer, 2003).

[33] Spearman, C. The proof and measurement of association between two things. Am. J. Psychol. 15, 72–101 (1904).

[34] Zimmerman, D. W. & Williams, R. H. The theory of test validity and correlated errors of measurement. J. Math. Psychol. 16, 135–152 (1977).

[35] Charles, E. P. The correction for attenuation due to measurement error: clarifying concepts and creating confidence sets. Psychol. Methods 10, 206–226 (2005).

[36] Campbell, D. T. & Fiske, D. W. Convergent and discriminant validation by the multitrait–multimethod matrix. Psychol. Bull. 56, 81–105 (1959).

[37] Richards, S. et al. Standards and guidelines for the interpretation of sequence variants. Genet. Med. 17, 405–424 (2015).

[38] Brnich, S. E. et al. Recommendations for application of the functional evidence PS3/BS3 criterion. Genome Med. 12, 3 (2020).

[39] Starita, L. M. et al. Variant interpretation: functional assays to the rescue. Am. J. Hum. Genet. 101, 315–325 (2017).

[40] McEwen, A. E. et al. Multiplexed assays of variant effect for clinical variant interpretation. Nat. Rev. Genet. 27, 137–154 (2025).

[41] Mann, H. B. & Whitney, D. R. On a test of whether one of two random variables is stochastically larger than the other. Ann. Math. Stat. 18, 50–60 (1947).

[42] Hardy, G. H., Littlewood, J. E. & Pólya, G. Inequalities 2nd edn (Cambridge Univ. Press, 1952).

[43] Esposito, D. et al. MaveDB: an open-source platform to distribute and interpret data from multiplexed assays of variant effect. Genome Biol. 20, 223 (2019).

[44] Rubin, A. F. et al. MaveDB 2024: a curated community database with over seven million variant effects from multiplexed functional assays. Genome Biol. 26 (2025).

[45] Efron, B. Nonparametric estimates of standard error: the jackknife, the bootstrap and other methods. Biometrika 68, 589–599 (1981).

[46] Tsuboyama, K. et al. Mega-scale experimental analysis of protein folding stability in biology and design. Nature 620, 434–444 (2023).

[47] Holm, S. A simple sequentially rejective multiple test procedure. Scand. J. Stat. 6, 65–70 (1979).

[48] Blackwell, D. Equivalent comparisons of experiments. Ann. Math. Stat. 24, 265–272 (1953).

[49] Lin, Z. et al. Evolutionary-scale prediction of atomic-level protein structure with a language model. Science 379, 1123–1130 (2023).

[50] Claussnitzer, M. et al. Minimum information and guidelines for reporting a multiplexed assay of variant effect. Genome Biol. 25 (2024).

